# Impact of Single Freeze-Thaw Cycles on Serum Protein Stability: Implications for Clinical Biomarker Validation Using Mass Spectrometry

**DOI:** 10.1101/2025.04.09.647925

**Authors:** T. Sauer, M. Oberländer, R. Maushagen, H. Radloff, R. Braun, K. Honselmann, U. Wellner, J. Habermann, C. Benecke, T. Keck, S. Szymczak, T. Gemoll

## Abstract

**Background:** Validation of biomarkers for clinical diagnostics and therapy necessitates the availability of a substantial number of high-quality samples, along with a complete set of clinical data. Establishing standards for sample collection, storage, and quality control is essential to reduce the variability of sample quality.

**Methods:** This study evaluated the impact of a single freeze-and-thaw (FT) cycle on the quality of liquid nitrogen-stored clinical serum aliquots. Data-independent acquisition mass spectrometry (DIA-MS) was performed to measure serum protein abundance in a test cohort comprising 25 patients and 99 samples, and a validation cohort comprising 109 patients/samples. Abundance differences of paired fresh and FT samples were assessed by employing biostatistics and bioinformatics approaches, including clustering analyses, linear mixed models, machine learning, and functional annotation.

**Results:** Following the library-free data analysis and implementation of strict data preprocessing procedures, data on the relative abundance of 213 (test cohort) and 248 (validation cohort) human serum proteins was available. Overall, the proteomic data demonstrated high quality and reproducibility. Significant changes in abundance between fresh and FT samples was observed for 30 proteins in the test cohort (q-value ≤0.05). Of these, 11 proteins (36.6%) were successfully validated in the validation cohort, including CP and IGHV6-1 which were identified as potential biomarker candidates for discriminating between patients with malignant or benign pancreatic disease. A correlation analysis of the measured protein intensity and storage duration revealed no significant association between the two variables.

**Conclusions:** Distinct protein abundance patterns can be discerned between fresh serum samples and samples stored in liquid nitrogen. The findings of this study suggest that FT cycles are an important pre-analytical factor and should be addressed during the translation of protein-based biomarkers into clinical use.

## Background

Cancer is one of the leading global health challenges, responsible for nearly 10 million deaths in 2020 alone, and is projected to become the leading cause of mortality by 2060 (1). The increasing incidence of various cancers, driven by factors such as lifestyle choices, environmental exposures, genetic predisposition, and an ageing population, underscores the need for effective diagnostic and therapeutic strategies. Liquid biopsies and protein-based biomarkers have emerged as promising tools in the fight against cancer, offering the potential for non-invasive detection and monitoring of tumor dynamics (2, 3). However, despite identifying numerous biomarker candidates through comprehensive basic research, the translation of promising molecules into clinical decision-making tools remains a challenge. Many candidates fail to demonstrate the required sensitivity and specificity in diverse populations, leading to a significant gap between discovery and application in real-world settings. There are many pitfalls in translating biomarkers: inadequate study design, study execution, and technical failures already during initial biomarker discovery (4). Interestingly, pre-analytical factors with an impact on sample quality in discovery and validation studies are often overlooked. Due to the need for large sample sizes, researchers typically rely on samples stored in biobanks, as it is not feasible to acquire sufficiently large numbers of fresh patient and control samples without transport or storage. Biobanks collect and store patient samples under strict regulations and, therefore, have high-quality samples and patient metadata at their disposal. However, in the clinical setting, samples are often analyzed under fresh conditions, without storage-related freeze-thaw (FT) cycles as a pre-analytical impact factor. This analyte discrepancy may be responsible for some challenges in the translation of promising biomarkers to clinical use.

Against this background, this study was conducted to comprehensively analyze the storage-related FT cycle as a pre-analytical impact factor influencing the global proteomic level of cancer patient serum samples using data-independent acquisition mass spectrometry (DIA-MS). Paired fresh and FT serum sample proteome profiles were compared to reveal distinct protein abundance patterns between both sample conditions. Furthermore, machine learning was applied to identify potential biomarkers among proteins that show FT sensitivity.

## Material and Methods

The study concept and workflow are visualized in **Figure 1A**.

**Figure 1:**
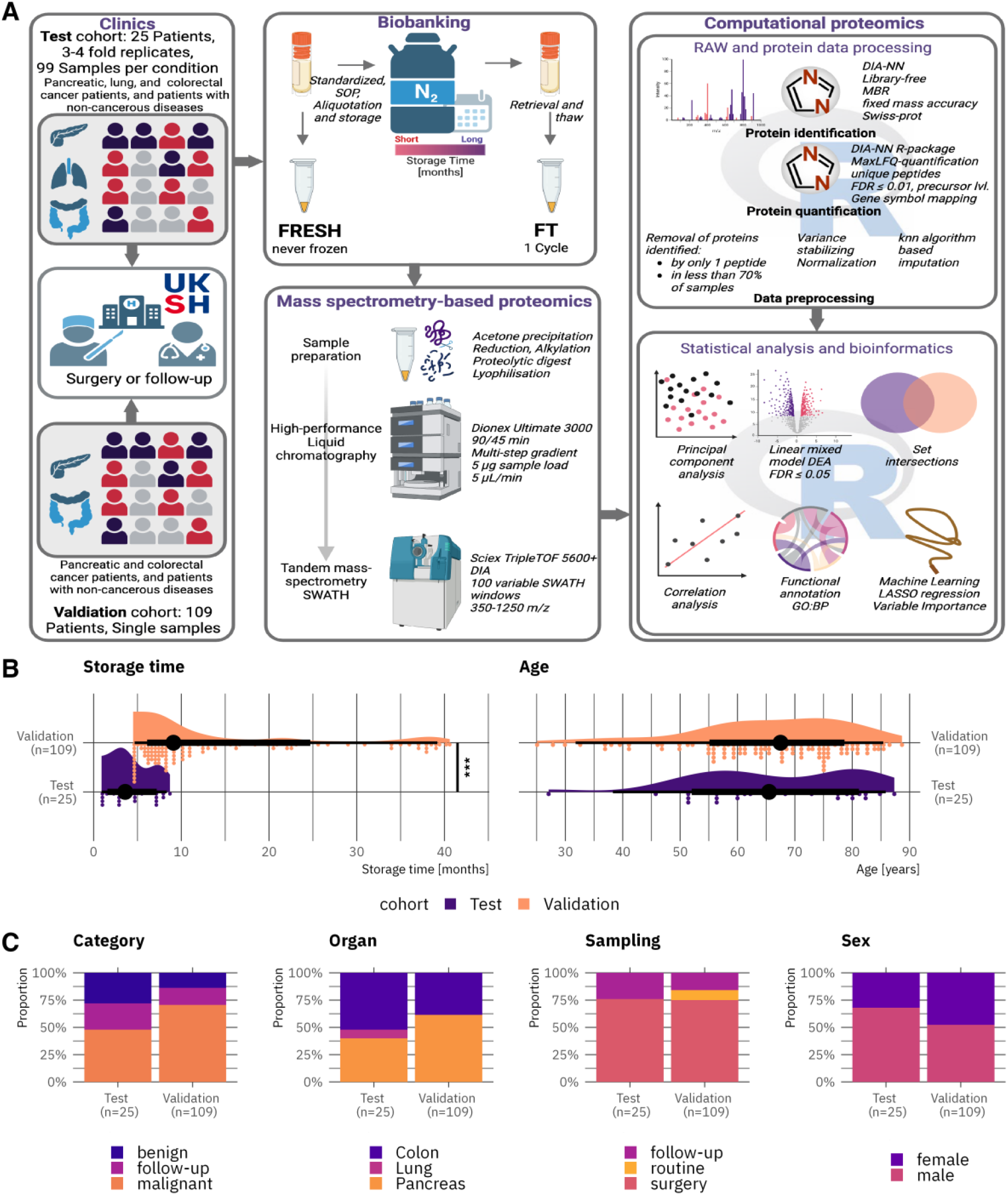
Study workflow and patient cohort meta data visualization. A: Study workflow description. Blood samples from two independent cohorts were collected in the clinic. Serum was derived from blood samples in the biobank and sample aliquots were stored in liquid nitrogen. Fresh aliquots (never frozen) were immediately processed and prepared for mass-spectrometry-based proteomics. FT samples (one freeze-and-thaw cycle) were extracted after a certain storage time and identically processed. Fresh and FT samples were analyzed with LC-MS/MS in data-independent acquisition mode. Acquired RAW data were processed in DIA-NN and the protein data were stringently preprocessed in R. Statistical and bioinformatics analyses were carried out in R and include clustering analyses, differential abundance analyses, correlation analyses, and functional annotations. A sub-cohort was used for a biomarker discovery simulation using machine learning. Asterisks indicate significance in the Wilcoxon rank sum test: *** = p-value ≤ 0.001. B: Raincloud plots visualizing the distribution of storage time (months) and patient age (years) across both study cohorts. C: Stacked bar plots representing meta data variables across both study cohorts.

### Overview of patient cohorts

Serum samples utilized in this study were sourced from samples collected for routine proteomic profiling at the Section of Translational Surgical Oncology and Biobanking, Department of Surgery, University Hospital Schleswig-Holstein, Campus Lübeck, Germany. Samples were obtained from patients prior to surgical intervention, during routine check-up appointments, or at follow-up visits. No exclusion criteria were applied to sample selection for this study. The research protocols were approved by the local ethics committees of the University of Lübeck (#16-281, #16-282, #17-043, and #19-147A, #2023-362). All participating patients provided informed written consent. Comprehensive patient characteristics are detailed in **Supplementary Table 1, Additional File 1**. Patients were categorized as either benign or malignant (‘category’ variable) based on their diagnosed condition. Patients whose blood samples were collected during post-surgical follow-up appointments were classified as ‘follow-up’ in the category and the ‘sampling’ variable. The ‘sampling’ variable further specifies the context of blood sample acquisition: ‘surgery’ denotes samples collected immediately before tumor removal surgeries, while ‘routine’ indicates samples collected during routine monitoring consultations within the Department of Surgery (without surgery).

### Test study cohort

The test study cohort consisted of three to four serum samples from 25 patients (acquired simultaneously), including 18 patients diagnosed with tumors (pancreatic, colorectal, and lung) and seven patients with non-neoplastic conditions. A total of 99 fresh and frozen-thawed (FT) serum sample pairs were analyzed in the test study. The median storage duration for the FT samples was 3.6 months, with a range of 0.9 to 8.7 months.

### Validation study cohort

The validation study utilized single measurements of 109 fresh and FT serum sample pairs. This cohort included serum samples from 94 patients with malignant pancreatic and colorectal cancers, as well as 15 patients with non-malignant pancreatic and colorectal conditions. The median storage duration of the FT samples within the validation cohort was 9.1 months, ranging from 4.5 to 40.6 months. The test and validation cohort were compared using the R *crosstable* package (v0.8.1) (5).

### Biobanking and Mass spectrometry-based proteomics

#### Serum sample acquisition

Following the acquisition of signed informed consent, peripheral blood was collected by the standard operating procedures established at the hospital-based Interdisciplinary Center for Biobanking Lübeck (University Hospital Schleswig-Holstein, Campus Lübeck, Germany). Patient blood samples were collected in 9 mL S-Monovettes Serum Gel (Sarstedt, Germany), allowed to cool at room temperature for 30 minutes, and subsequently centrifuged at 1,500 × g, 10 min and 4°C. The serum was then aliquoted into 0.7 mL cryovials (Azenta Life Sciences, USA). Before freezing in liquid nitrogen, a 10 μL sub-aliquot of serum was transferred to 1.5 mL low protein binding tubes and stored on ice during transport to the Section of Translational Surgical Oncology and Biobanking laboratories for immediate sample preparation and protein digestion.

#### Protein digestion

For FT sample preparation, stored serum samples were retrieved from liquid nitrogen. A 2 μL aliquot of serum was combined with 18 μL milli-Q/ultra-pure (MQ) water. Subsequently, 10 μL of the diluted serum was added to 40 μL of ice-cold 100% acetone, vortexed, and incubated at -20°C for 1h. The serum was then centrifuged for 10 min at 10,000 × g and 4°C. The supernatant was carefully removed and discarded, while the resulting pellet was allowed to air-dry. The pellet was resuspended in 100 μL deoxycholate (DOC) buffer (7.9 mg ammonium bicarbonate, 1 mL milli-Q water, and 10 mg sodium deoxycholate) and 1 μL 1,4-dithiothreitol 1M solution in DOC buffer. This solution was incubated at 60°C for 30 min. Subsequently, 4 μL of 0.5M iodoacetamide was added and incubated for 30 min in the dark. Afterwards, 1 μL of trypsin was added, and the mixture was incubated overnight at 37°C. The digestion was quenched with the addition of 1 μL of formic acid, and the samples were centrifuged for 5 min at 14,000 × g. The resulting supernatant was transferred to a new tube and frozen at -80°C. Thereafter, the samples were lyophilized for 3h in a vacuum centrifuge (Christ Gefriertrocknungsanlagen GmbH, Germany). The serum digest was then stored at -20°C until further use.

#### High-performance liquid chromatography

The serum sample digests were solubilized in 50 mL loading buffer and separated using a Dionex Ultimate 3000 high-performance liquid chromatography (HPLC) system (Thermo Fisher, USA). The HPLC was configured with a precolumn (μ-Precolumn Acclaim PepMap100, 0.3 mm × 5 mm, 5 μm particle size, 100 Å pore size, Thermo Fisher Scientific, USA). A Luna C18(2) (0.3 mm × 50 mm, 3 μm particle size, 100 Å pore size, Phenomenex Inc., USA) was used as an analytical column. Although the identical chromatographic system was employed for both study cohorts, the gradient settings differed between the test and validation studies. A 90-minute gradient was employed to analyse the test study cohort serum samples. The method commenced with a 4 min sample injection and trap column loading, utilizing an initial solvent composition of 97% solvent A (1% formic acid in MQ water) and 3% solvent B (1% formic acid in acetonitrile). Subsequently, a linear gradient was applied, increasing the proportion of solvent B to 25% at 68 min, 35% at 73 min, and finally reaching 80% at 80 min, which was sustained for 3 min. Subsequently, the concentration of solvent B was gradually reduced to 3% over a 4 min period, which was maintained for 8 min.

The validation study employed a 45-minute gradient, proportionally shortened to the 90-minute gradient used in the test study, to increase the sample throughput. Accordingly, the 45 min gradient profile comprised a 4 min sample injection and trap column loading at 97% solvent B to 3% solvent A, followed by a linear gradient, increasing the proportion of solvent B to 25% at 30.68 min, 35% at 32.94 min, and 80% at 33.84 min until 35.2 min. Subsequently, the proportion of solvent B was reduced to 3% at 37 minutes and maintained for 8 min. A flow rate of 5 μl/min was used for all HPLC methods.

#### Mass spectrometry

Peptides eluting from the Dionex Ultimate 3000 HPLC were analyzed using a TripleTOF 5600+ (AB Sciex, USA) mass spectrometer system. For serum sample data acquisition, data-independent acquisition-MS in sequential window acquisition of all theoretical mass spectra (SWATH) was employed. The following acquisition parameters were used: Ion Spray Voltage Floating at 5000 V; ion source gas (GS1), 15; ion source gas (GS2), 0; curtain gas at 30 and source temperature heating set to 0 °C. The optimized declustering potential was set at 100; collision energy to 19.2; collision energy spread, 5.0; ion release delay, 67; ion release width at 25. One 0.05 s MS scan (m/z 350–1250) was performed for data acquisition. Based on the 90- and 45-minute HPLC gradients, two 100 precursor isolation window SWATH methods were developed. The precursor isolation windows were defined using the SWATH Variable Window Calculator V1.1 (AB Sciex, USA) based on precursor *m*/*z* densities obtained from data-dependent acquisition (DDA) spectra. The mass range was set to 350-1250 m/z, window overlap was set to 1.0 Da, and collision energy spread was fixed at 5.0 eV. For DDA acquisition, identical instrument working parameters were used. MS scans were performed for 350-1250 m/z with an accumulation time of 0.25 sec, and MS/MS scans were performed for 100 – 1500 m/z with an accumulation time of 0.05 s at high sensitivity mode.

### Computational proteomics

#### SWATH data processing

The raw SWATH data from both study sub-cohorts were processed using the software tool *DIA-NN* v1.8.1 with identical settings (6). The software was used in the high-accuracy LC mode with retention time-dependent cross-normalization enabled. Mass accuracy was fixed to 2e-05 (MS2) and 1.5e-05 (MS1), while scan window width was determined automatically. The ‘match between runs’ function was used to develop a spectral library using the ‘smart profiling strategy’ from the data-independent acquisition data. The human UniProtKB/swiss-prot database (version 2020/12/6) (7) was used for protein inference from identified peptides. Trypsin/P was specified as protease. The precursor ion generation settings were set to a peptide length of 7–52 amino acids, the maximum number of missed cleavages to one. The maximum number of variable modifications was set to zero. N-terminal methionine excision and cysteine carbamidomethylation were enabled as fixed modifications.

### Statistical analysis and bioinformatics

#### Protein quantity data preprocessing

The DIA-NN report files were further processed in the *DIA-NN R* package (v1.0.1) (6) for MaxLFQ-based protein quantification (8). For the test and validation dataset, a report was generated containing the unique proteins that passed the FDR cut-off of 0.01 applied at the precursor level and were identified and quantified using proteotypic peptides only. A minimum of two unique peptides identified all proteins included in the final datasets. The proteins were mapped to their corresponding gene names; thus, the terms ‘proteins’ and ‘genes’ are used interchangeably in this study. The quantitative data were further pre-processed using the R package *DEP* (v1.26.0) (9). The unique protein data sets were filtered for data completeness: each protein had to be identified in at least 70% of samples. A variance stabilizing normalization (10) was applied to the data, including a log^2^ transformation, before imputing the remaining missing values using the k-nearest neighbour model. DEP borrows the imputation function from *MSnbase* and was used with default settings *(11, 12)*.

#### Proteomic data quality assessment

The two acquired proteomic datasets were assessed on their reproducibility across the acquisition periods by evaluating the protein and peptide identifications, normalized intensity ranges, and coefficients of variation (CV) across acquisition batches. Differences within CV was assessed utilizing an ANOVA and/or Student’s t-test, corrected for multiple testing with the Benjamini-Hochberg procedure (13), using the *ggpubr* package (v0.6.0) (14).

#### Assessment of FT cycle impact

A linear mixed model (LMM) was used to perform a two-group comparison of paired fresh and FT sera for each protein separately, employing the *nlme* R-package (v3.1-167) (15). Protein abundance was modelled as dependent variable with type of sample (fresh or FT) as fixed and the subject ID as random effects. The Benjamini-Hochberg procedure was used to adjust p-values for sample type (13). A significant difference in protein levels between fresh and FT samples was considered at q < 0.05. Principal component analyses (PCA) were calculated for both study cohorts using the base R *stats* package (v4.4.2) (16). Correlation analysis was carried out using the *ggpubr* package (v0.6.0) (14) and the R package *corrplot (v0*.*95)* (17) utilizing the protein intensity data of the validation cohort. The *tidyverse* package (v2.0.0) collection was employed throughout all bioinformatics analyses to facilitate data handling and visualization (18). Key packages used include *dplyr (v1*.*1*.*4)* for data manipulation and *ggplot2 (v3*.*5*.*1)* for visualization.

#### Physiochemical property mapping

For validated target proteins, the physiochemical properties hydrophobicity index (Kyte & Doolittle scale) (19), the instability index (20), and the aliphatic index (21) were calculated using the Peptides R package developed by Osorio et al. (22). The complete amino acid sequences of the proteins were extracted from UniProtKB/swiss-prot database (version 2024/10/28) and used to calculate the parameters. The variables hydrophobic index, instability index, and aliphatic index were discretized for better visualization as follows: hydrophobicity: >0 ≙hydrophobic,<0 ≙hydrophilic; instability index: >40 ≙unstable, <40 ≙stable; aliphatic index: <70 ≙low, 70-90 ≙moderate, >90 ≙high.

#### Identification of biomarker candidates using machine learning

Fresh serum proteomics from a sub-cohort of 46 patients within the validation cohort, encompassing individuals with pancreatic ductal adenocarcinoma (PDAC) (n = 27) and benign lesions (n = 10), were analysed to identify potential biomarkers. Preprocessing included the removal of zero-variance predictors and the z-transformation of all predictor variables. A dataset split was applied to the data, with 80% of the data allocated to the training and 20% to the testing set. Stratification was performed on the target variable (PDAC *vs*. Benign).

A Least Absolute Shrinkage and Selection Operator (LASSO) regression model was employed to discern key features for classifying samples into benign and malignant categories using the *tidymodels (v1*.*3*.*0)* framework (23). The penalty hyperparameter was tuned by 10-fold cross-validation across a grid of penalty values ranging between 1e^-10^ and 1e^10^. To enhance feature selection, the 10^th^ best lambda value was selected based on the root mean squared error RMSE. This less stringent approach permitted the inclusion of additional features in the final model. The final LASSO model was then fitted to the training data, and non-zero coefficients (feature importances) were extracted to identify the selected features and their impact on the target variable in terms of classification in the malignant category.

### Data availability

The mass spectrometry data have been deposited at the ProteomeXchange Consortium via the PRIDE partner (24) repository with the dataset identifier PXD060473.

## Results

### Study workflow and cohort description

This study investigated the pre-analytical effects of a single freeze-thaw (FT) cycle on the serum proteome, specifically within the context of cancer biomarker research (**Figure 1A**). A paired comparison was conducted, analyzing serum samples in their fresh state (never frozen) and after undergoing a single FT cycle following storage in liquid nitrogen for varying durations. The study employed a test cohort of 99 fresh-FT sample pairs from 25 patients and a validation cohort of 109 fresh-FT sample pairs derived from 109 patients. The samples were subjected to mass spectrometry-based proteomics in data-independent acquisition. Computational proteomics, incorporating rigorous data pre-processing of the RAW and protein data, statistical analysis, and bioinformatics, was subsequently performed. This analysis aimed to evaluate protein profiles of fresh and FT serum samples and to characterize the impact of an FT cycle on the proteome. The descriptive statistical analysis results are presented in **Table 1**, while **Figure 1B-C** illustrates the variable distribution across both cohorts. The median storage duration for the test cohort was 3.6 months, while the validation cohort demonstrated a median storage time of 9.1 months. The diseased organ distribution was different between the cohorts. Specifically, the validation cohort comprises a higher proportion of patients with colon diseases, while pancreatic diseases were predominant within the validation cohort.

**Table 1:**
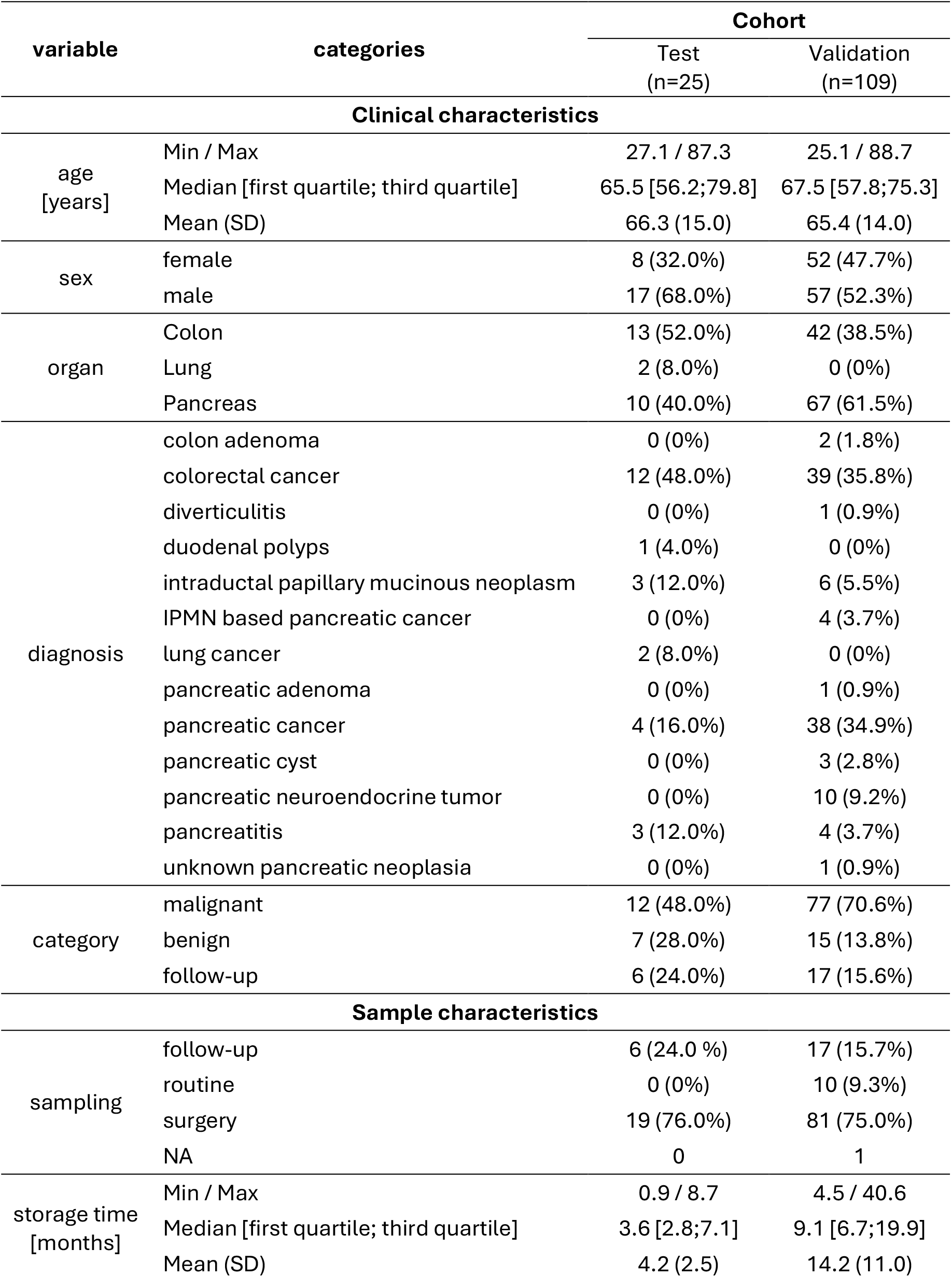
Comparison of study cohorts based on clinical and sample characteristics.

| variable | categories | Cohort |  |
| --- | --- | --- | --- |
|  |  | Test<br>(n=25) | Validation<br>(n=109) |
| Clinical characteristics |  |  |  |
| age<br>[years] | Min / Max | 27.1 / 87.3 | 25.1 / 88.7 |
|  | Median [first quartile; third quartile] | 65.5 [56.2;79.8] | 67.5 [57.8;75.3] |
|  | Mean (SD) | 66.3 (15.0) | 65.4 (14.0) |
| sex | female | 8 (32.0%) | 52 (47.7%) |
|  | male | 17 (68.0%) | 57 (52.3%) |
| organ | Colon | 13 (52.0%) | 42 (38.5%) |
|  | Lung | 2 (8.0%) | 0 (0%) |
|  | Pancreas | 10 (40.0%) | 67 (61.5%) |
| diagnosis | colon adenoma | 0 (0%) | 2 (1.8%) |
|  | colorectal cancer | 12 (48.0%) | 39 (35.8%) |
|  | diverticulitis | 0 (0%) | 1 (0.9%) |
|  | duodenal polyps | 1 (4.0%) | 0 (0%) |
|  | intraductal papillary mucinous neoplasm | 3 (12.0%) | 6 (5.5%) |
|  | IPMN based pancreatic cancer | 0 (0%) | 4 (3.7%) |
|  | lung cancer | 2 (8.0%) | 0 (0%) |
|  | pancreatic adenoma | 0 (0%) | 1 (0.9%) |
|  | pancreatic cancer | 4 (16.0%) | 38 (34.9%) |
|  | pancreatic cyst | 0 (0%) | 3 (2.8%) |
|  | pancreatic neuroendocrine tumor | 0 (0%) | 10 (9.2%) |
|  | pancreatitis | 3 (12.0%) | 4 (3.7%) |
|  | unknown pancreatic neoplasia | 0 (0%) | 1 (0.9%) |
| category | malignant | 12 (48.0%) | 77 (70.6%) |
|  | benign | 7 (28.0%) | 15 (13.8%) |
|  | follow-up | 6 (24.0%) | 17 (15.6%) |
| Sample characteristics |  |  |  |
| sampling | follow-up | 6 (24.0 %) | 17 (15.7%) |
|  | routine | 0 (0%) | 10 (9.3%) |
|  | surgery | 19 (76.0%) | 81 (75.0%) |
|  | NA | 0 | 1 |
| storage time<br>[months] | Min / Max | 0.9 / 8.7 | 4.5 / 40.6 |
|  | Median [first quartile; third quartile] | 3.6 [2.8;7.1] | 9.1 [6.7;19.9] |
|  | Mean (SD) | 4.2 (2.5) | 14.2 (11.0) |

### Proteomics data quality and comparability

Serum proteomic profiles were generated for both study cohorts using a library-free data analysis approach. The test study resulted in a proteomic dataset of 198 samples, acquired across five batches over a period of approximately 43 weeks. The validation cohort data encompassed 218 samples, distributed across six batches over approximately 60 weeks. Protein identification rates remained consistent for both proteomic datasets throughout the acquisition period and across all acquisition batches. The test cohort analysis identified between 203 and 230 unique proteins (mean 214.7, standard deviation 5.08), while the validation cohort analysis identified between 236 and 271 unique proteins (mean 256.5, standard deviation 6.43, **Figure 2A&D**). Following pre-processing, the datasets contain relative abundance data for 213 unique proteins in the test cohort and 248 unique proteins in the validation cohort. The number of identified peptides per protein was consistent across both cohorts, with a mean of eight peptides per protein (**Figure 2B&E**). A statistically significant difference in mean coefficient of variation (CV) between batches was identified exclusively within the test cohort (Anova p-value = 0.028), while no such difference was evident in the validation cohort (p-value = 0.7) (**Figure 2C&F)**. Pairwise comparisons between test study batches revealed a significant difference in the mean coefficient of variation for P.A *versus* P.D, and P.E, respectively (Student’s t-test adj.p ≤ 0.05). However, these results suggest a high degree of reproducibility and inter-batch comparability; therefore, no batch correction was applied to the datasets.

**Figure 2:**
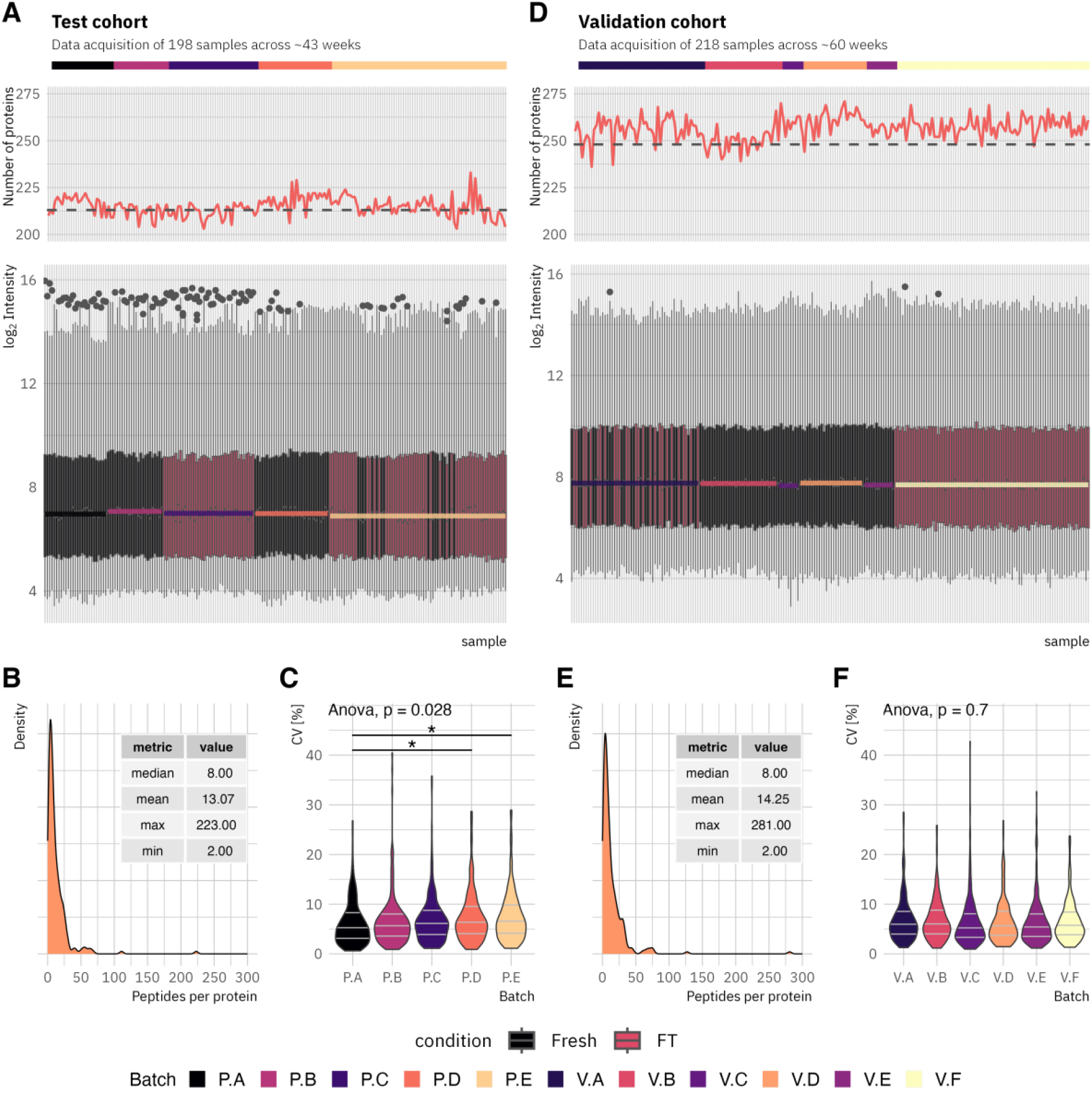
Proteomics data quality control for test **(A-C)** and validation cohort data **(D-F)**. The test cohort data were acquired in five batches over approximately 43 weeks. The validation cohort samples were measured in six batches over approximately 60 weeks. **A&D:** Visualization of the normalized log_2_ intensity for all samples in chronological order as box plots, the number of proteins identified per sample before filtering as line plot (dashed line indicates the number of proteins in the final, pre-processed dataset), as well as the corresponding acquisition batch per sample. Colored lines in boxplots represent the average median of the corresponding batch. **B&E:** Density plots visualizing the distribution of identified peptides per protein, including descriptive metrics as table. **C&F**: Violin plots visualizing the distribution of the CV per analysis batch. Grey lines within violins indicate quantiles. Asterisks indicate significance in students’ t-test: * = p-value ≤ 0.05, ** = p-value ≤ 0.01.

### FT cycle as pre-analytical impact factor

A principal component analysis was performed on both proteomic datasets to assess global disparities in proteomic profiles. The analysis revealed a relatively ambiguous clustering of FT and fresh samples of the test cohort, largely influenced by the high similarity of sample replicates (**Figure 3A**). In contrast, the validation cohort exhibited a distinct clustering trend, particularly captured by PC2 (6.1% total explained variance).

**Figure 3:**
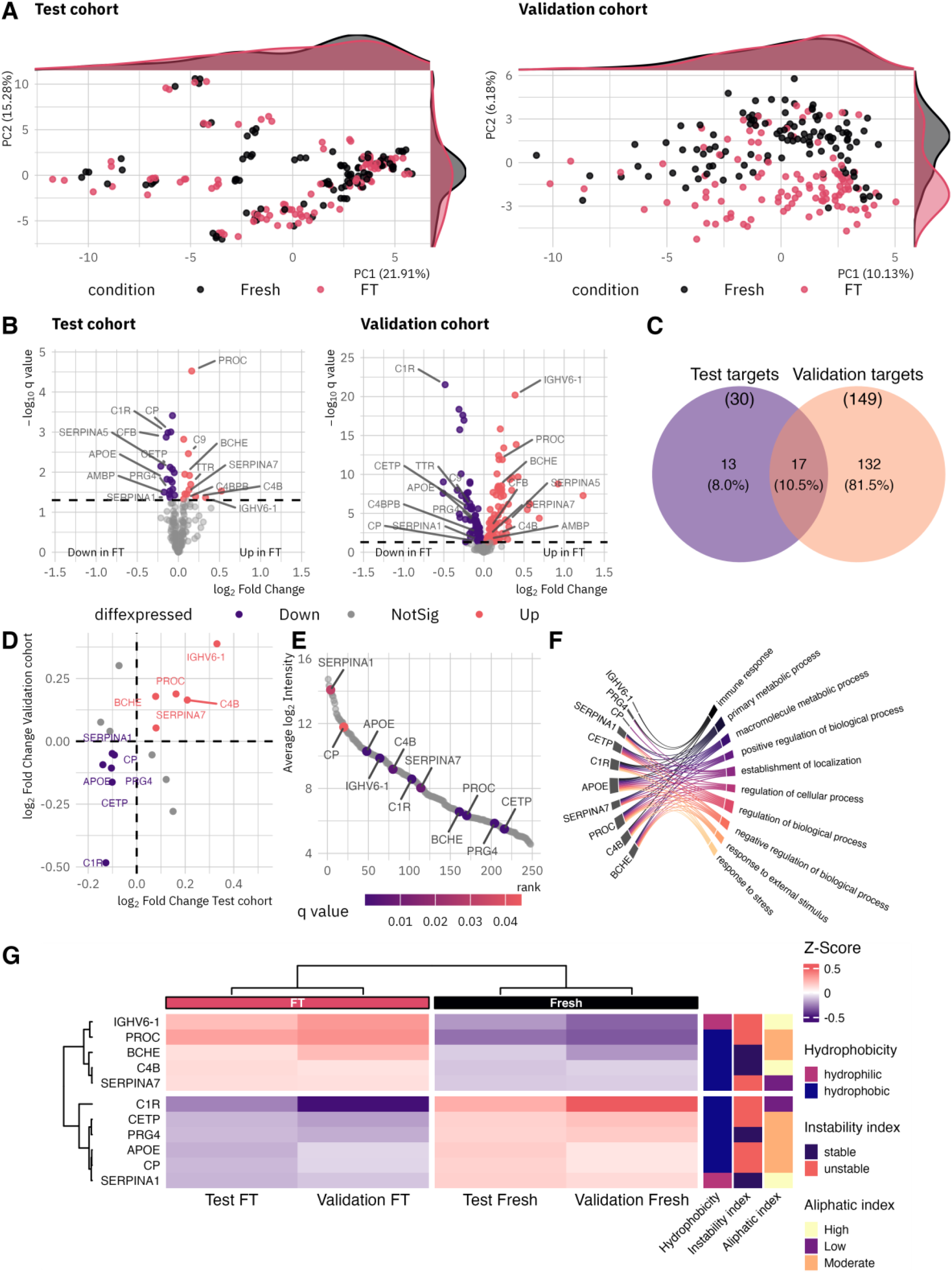
Proteomic profiling results for the test and validation cohorts. **A:** Principal component analysis plots for the test and validation cohorts. While the clustering of the test cohort is relatively indistinct, the clustering of the validation cohort shows clear tendencies. Adjacent density plots show the sample group overlaps by showing the kernel densities. Percentages of total variance explained by each component are shown in parentheses. **B:** Volcano plots visualize the results of differential intensity analysis using LMMs. Significance is considered at q-values ≤ 0.05. While 30 proteins were found to be differentially abundant in the test cohort, 149 proteins showed significant differences in abundance in the validation cohort. Protein names highlight proteins identified as differentially abundant in both cohorts. **C:** Venn diagram visualizing the total overlap of significant proteins identified in the test and validation studies. **D:** Scatterplot visualizing the fold change of each protein in test and validation cohort. **E:** Protein rank plot from highest to lowest abundant proteins (calculated from the validation cohort), illustrating the dynamic range of target proteins. **F:** Chord plot visualizing the top 10 gene ontology mappings for the biological process subontology, selected by gene ratio. **G:** Heatmap illustrating average Z-score transformed protein intensities for target proteins, calculated by cohort and condition (fresh and FT). Heatmap annotations illustrate important physiochemical properties: Kyte & Doolittle hydrophobicity scale (discretized: >0 ≙hydrophobic, <0 ≙hydrophilic), Guruprasad instability index (discretized: >40 ≙unstable, < 40 ≙stable), and Ikai aliphatic index (discretized: <70 ≙low, 70-90 ≙moderate, >90 ≙high).

To identify proteins exhibiting significant intensity variations in FT samples, linear mixed models were applied to the proteomic profiles of both the test and validation cohorts. The analysis revealed statistically significant differential intensity (q-value ≤ 0.05) for 30 proteins when comparing paired fresh and FT samples of the test cohort (**Figure 3B**). Specifically, 17 proteins (57%) demonstrated increased intensity, while 13 (43%) exhibited decreased intensity in FT samples (detailed results for the test cohort are available in **Supplementary Table 2, Additional File 1**). Overall, 14% of the identified serum proteins demonstrated a statistically significant difference in measurable intensity following one FT cycle. The validation study further corroborated these findings, with 149 proteins (60%) demonstrating a statistically significant alteration in measured intensity following one FT cycle due to storage in liquid nitrogen (**Figure 3B**). Notably, the proportion of proteins exhibiting increased or decreased intensities in FT samples was consistent with the test study. Specifically, 70 (47%) displayed an increased fold-change, while 79 proteins (53%) showed decreased intensities (detailed results for the validation cohort are available in **Supplementary Table 3, Additional File 1**). A Venn diagram, depicted in **Figure 2C**, illustrates the intersection of differentially expressed proteins. Among the 162 unique proteins identified as FT-sensitive, 17 were observed in both the test and validation cohorts. Of these 17 proteins, 11 proteins exhibited consistent directional changes (either decreased or increased intensity) across both cohorts (**Figure 3D**). These eleven proteins were designated as validated FT-sensitive proteins and spanned approximately eight orders of magnitude (**Figure 3E**). Functional annotation by mapping the gene symbols to the subontology ‘biological process’ of the Gene Ontology (GO) database revealed involvement in the immune response, metabolic processes, cellular and biological processes, the establishment of localization, and the response to stress (**Figure 3F**). Detailed GO:BP mapping results are available in **Supplementary Table 4, Additional File 1**.

A data mining approach was applied to assess the physiochemical attributes of the target proteins (22). In this context, three characteristics were identified as particularly relevant: hydrophobicity index, instability index, and aliphatic index. All indices were calculated based on the primary amino acid sequence of each protein. **Figure 4G** depicts the protein annotation results, including the mean relative protein intensity by condition and cohort. Of the eleven proteins analyzed, only two showed hydrophilic characteristics (index >0 ≙hydrophobic, <0 ≙hydrophilic). Seven of the eleven presented an instability index >40, indicating instability (index >40 ≙unstable, <40 ≙stable). Two proteins demonstrated a low aliphatic index, six were moderate, and three had a high aliphatic index (index <70 ≙low, 70-90 ≙moderate, >90 ≙high). Each characteristic phenotype was represented by proteins with either increased or decreased intensity following the FT cycle. The target protein data, along with the physiochemical annotation results, are available in **Supplementary Table 5, Additional File 1**.

**Figure 4:**
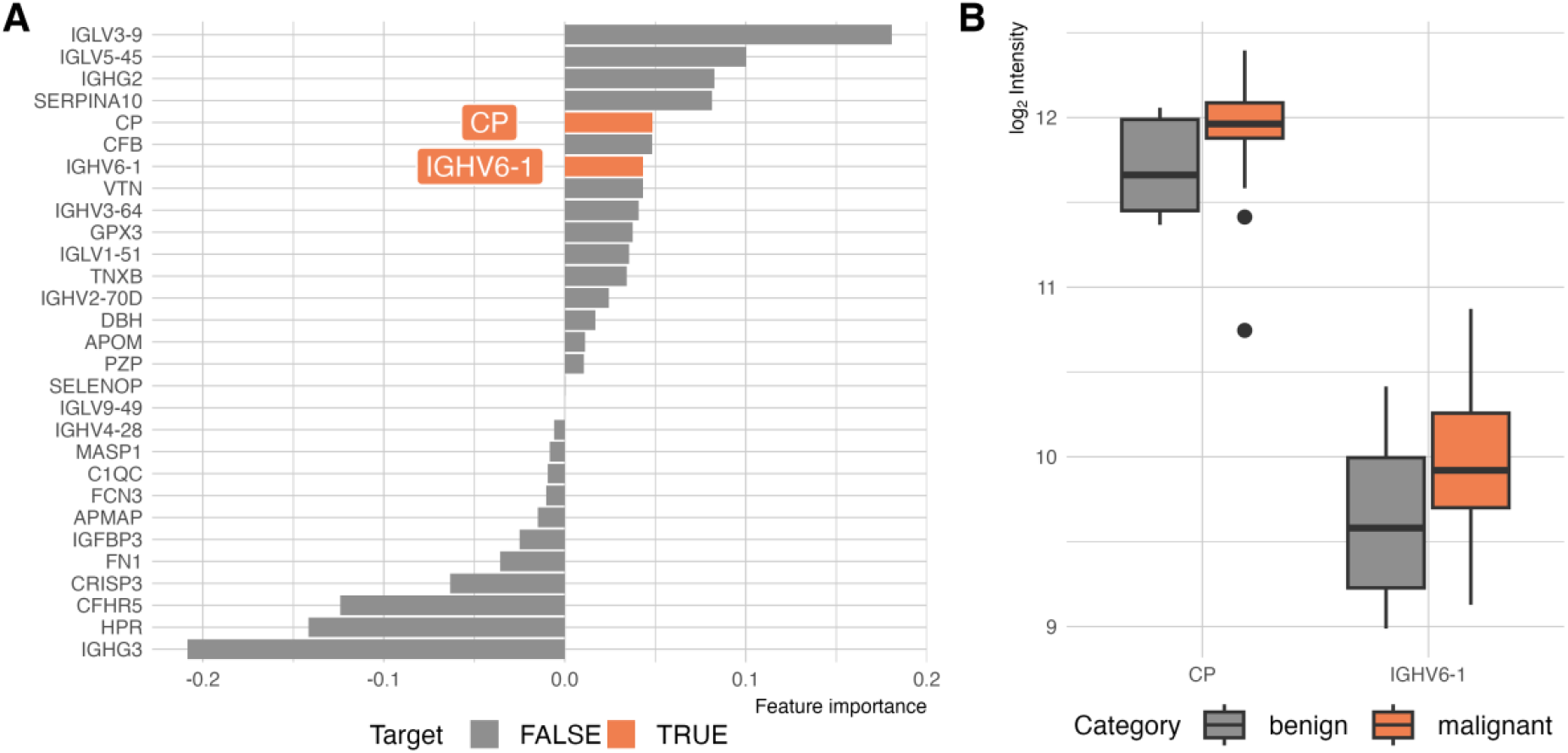
Biomarker candidate identification for classification into malignant and benign pancreatic diseases. A data subset from 37 fresh validation cohort serum samples was used (27 malignant, 10 benign). **A:** LASSO regression results used for feature selection. Two target proteins were among the 29 features with an estimate ≠ 0. **B:** Boxplots visualizing the measured log_2_ intensity of CP and IGHV6-1.

To evaluate the impact of the storage time in liquid nitrogen on quantifiable protein intensity within FT samples, a a analysis was performed using all FT samples from the validation cohort. The results indicated no significant correlation between storage time and protein intensity. The correlation analysis results are visualized in **Supplementary Figure 1, Additional File 2**. Complement component 1r (C1R) exhibited the highest net Pearson correlation coefficient (R = 0.22, p-value = 0.019). Furthermore, no strong correlation was identified between the storage time and patient age, with Cholinesterase (BCHE) showing the highest net R of -0.36 (p-value = 0.0001) (see **Supplementary Figure 1, Additional File 2**).

These findings suggest that a single FT cycle is sufficient to induce a sustainable effect on serum samples, which is quantifiable through both clustering and differential expression analyses. The duration of storage does not appear to influence the levels of FT-sensitive proteins.

### FT-sensitive proteins are potential biomarker candidates

The identification of FT-sensitive proteins as potential biomarker candidates presents a challenge for the translation of findings from academic research, where FT samples are common, to clinical applications, which typically utilize fresh samples. Consequently, fresh serum proteomics data from a sub-cohort of 37 patients within the validation cohort were analysed. This sub-cohort comprised patients with malignant pancreatic diseases (pancreatic ductal adenocarcinomas/PDAC, n = 27) and benign pancreatic lesions (n = 10). Details regarding sample inclusion are provided in **Supplementary Table 1, Additional File 1**. LASSO regression analysis identified 37 proteins as relevant for this classification problem, with non-zero estimated coefficients. The model achieved an accuracy of 0.62 and an area under the receiver operator characteristics curve of 0.67. Notably, two of these proteins were also identified as FT-sensitive: IGHV6-1 (Immunoglobulin heavy variable 6-1) and CP (Ceruloplasmin) (**Figure 4A**). Both proteins exhibited an increased median log^2^ intensity in malignant samples compared to benign samples (**Figure 4B**). These findings indicate that FT-sensitive proteins may be identified as proteomic biomarkers in serum samples highlighting the significance of FT-cycle-mediated factors in translational research.

## Discussion

The translation of biomarker candidates from basic research to clinical application remains a substantial challenge in modern medicine. While rapid advancements in omics technologies have facilitated the identification of numerous promising biomarker candidates for novel diagnostics and therapeutics, the translation process is hindered by obstacles such as a lack of standardized protocols for sample collection and processing, as well as insufficient quality control measures (4, 25, 26). These factors can result in the identification of biomarkers that lack the necessary sensitivity and specificity for clinical use. To address these challenges, our study aimed to quantitively assess the pre-analytical effect of a storage-related freeze-thaw (FT) cycle on the serum proteome. Using mass-spectrometry, an analytical technique uniquely suited for analyzing fresh samples, we examined over 200 fresh and FT serum sample pairs.

The proteomic data quality was rigorously evaluated throughout the duration of the study (43 weeks for the test and 60 weeks for validation cohort). Although the study design was not optimal, with some batches being confounded by sample type, overall, no significant batch effects were detected between the sample processing order, signal intensities, storage duration, or protein identifications, demonstrating the high reproducibility of the proteomic approach used and its suitability for large-scale quantitative proteomic profiling. In this context, clustering analysis of the test study data showed that sample replicates from the same patient were the primary determinant of clustering, further emphasizing the reproducibility of DIA-MS measurements. The validation study revealed a distinct clustering trend between fresh and FT samples, indicating a global and systemic effect of the FT cycle on the proteome. Despite of the observed low effect sizes, 11 out of 30 initially identified target protein candidates were validated across both studies and their FT sensitivity should be considered in biomarker studies. In line, these validated proteins spanned a broad dynamic range and were involved in various biological processes, indicating that FT cycle sensitivity is not limited to specific protein groups or abundance levels. Furthermore, the validated FT-sensitive proteins exhibited diverse physiochemical properties, including thermodynamically stable and unstable proteins. A majority were classified as unstable and hydrophobic. Indeed, the hydrophobic effect is recognized as a critical determinant in maintaining protein core stability during protein folding and cold denaturation (27-30). At lower temperatures, a reduction in the hydrophobic effect can lead to an increased tendency for cold denaturation of proteins (31). Therefore, hydrophobic proteins may be more susceptible to the effects of the FT cycle than their hydrophilic counterparts. Additionally, observed differential protein quantities after one FT-cycle likely result from forms of cryo-damage. Cryo-damage during freezing and thawing is mainly due to reversible protein aggregation caused by ice crystal growth (32-35), including cryoconcentration (36), contact with interfaces (37, 38), and cold denaturation (39, 40). While reduced protein abundance typically indicates damage, increased quantities may result from degradation and misfolding of protein complexes.

Another finding that stands out from the results pertains to the potential impact of sample storage conditions on biomarker detection in a cohort of 37 patients diagnosed with pancreatic disease (27 malignant, 10 benign). LASSO regression detected 29 features as important in distinguishing between benign and malignant samples. Notably, two of these proteins, IGHV6-1 and CP, were also identified as FT-sensitive, highlighting the need to evaluate pre-analytical impact factors during the translation of biomarkers in the future. Both showed higher median log^2^ intensities in malignant samples, suggesting their potential as biomarkers for malignant pancreatic cancer.

The multicopper oxidase, ceruloplasmin (CP) is primarily synthesized in the liver and subsequently released into the bloodstream (41). This multifunctional protein participates in iron and copper metabolism, antioxidant activity, and amine oxidation (42). Furthermore, CP functions as an acute-phase protein, responding to physiological stressors including inflammation and trauma (43). It has already been classified as a prognostic biomarker in various cancer entities, including breast cancer (44), glioma (45) and bile duct cancer (46). Interestingly, elevated serum CP levels have also been reported in patients with pancreatic cancer (47). Hanas et al. proposed that pancreatic cancer may exhibit a vulnerability to chemical agents that disrupt copper homeostasis, such as ammonium tetrathiomolybdate in the treatment of squamous cell carcinoma of the head and neck (48). IGHV6-1, the V region of the variable domain of immunoglobulin heavy chains, participates in antigen recognition (49) and has not been described as cancer biomarker candidate.

## Conclusions

This study highlights the importance of high-quality samples and standardized handling in biomarker research. Using data-independent acquisition mass spectrometry, it demonstrates the significant impact of freeze-thaw cycles on stored serum samples. This pre-analytical impact factor may explain why many proposed protein biomarkers fail to translate to clinical applications. The findings underscore the need for rigorous quality management in biospecimen handling and could improve biomarker research.

## List of abbreviations

BCHE: Cholinesterase
C1R: Complement component 1r
CP: Ceruloplasmin
CV: coefficient of variation
DDA: data-dependent acquisition
DIA-MS: Data-independent acquisition mass spectrometry
DOC: deoxycholate
FDR: false-discovery rate
FT: freeze-and-thaw
GO: gene ontology
HPLC: High-performance liquid chromatography
IGHV6-1: Immunoglobulin heavy variable 6-1
LASSO: least absolute shrinkage and selection operator
LMM: linear mixed model
MQ: ultra-pure water
MS: mass spectrometry, mass spectrometry
MS/MS: tandem mass-spectrometry
PCA: principal component analysis
PDAC: Pancreatic ductal adenocarcinoma
RMSE: root mean square error
SWATH: sequential window acquisition of all theoretical mass spectra

## Additional files

Additional_file_1.xlsx: Supplementary Table 1: Patient cohort clinical and meta data; Supplementary Table 2: Differential expression analysis results of the test cohort study; Supplementary Table 3: Differential expression analysis results of the validation cohort study; Supplementary Table 4: Gene ontology ‘Biological process’ mapping results of the validated target proteins; Supplementary Table 5: Physiochemical annotation data of the validated target proteins.

Additional_file_2.docx: Supplementary Figure 1: Correlation analysis results of target proteins and storage duration.

## Declarations

### Ethics approval and consent to participate

The studies were approved by the local ethics committees of the University of Lübeck, namely #16-281, #16-282, #17-043, and #19-147A. All patients provided informed written consent.

### Consent for publication

Not applicable.

### Availability of data and materials

The mass spectrometry proteomics data have been deposited to the ProteomeXchange Consortium via the PRIDE partner repository with the dataset identifier PXD060473.

### Competing interests

The authors declare that they have no competing interests.

### Funding

This study was performed as a part of the AKELOP project (project number LPW-E/1.2.2/673) supported by State Program Economy 2014-2020 with funds from the European Regional Development Fund (ERDF) and the state Schleswig-Holstein. Thorben Sauer received a doctoral degree scholarship from the Ad Infinitum foundation.

### Authors’ contributions

**TS:** Data curation (lead), Formal analysis (lead), Investigation (equal), Methodology (equal), Software (lead), Validation (lead), Visualization (lead), Writing – Original Draft Preparation (lead). **MO:** Resources (supporting), Writing – Review & Editing (equal). **RM:** Resources (supporting), Writing – Review & Editing (equal). **HR:** Data curation (supporting), Writing – Review & Editing (equal). **RB:** Resources, Writing – Review & Editing (equal). **KH:** Resources (supporting), Writing – Review & Editing (equal). **UW:** Resources (supporting), Writing – Review & Editing (equal). **JH:** Resources (supporting), Writing – Review & Editing (equal). **CB:** Resources (supporting), Writing – Review & Edition (equal). **TK:** Resources (supporting), Writing – Review & Editing (equal), **SS:** Methodology (equal), Data curation (supporting), Writing – Review & Editing (equal). **TG:** Funding Acquisition (lead), Project administration (lead), Resources (lead), Validation (supporting), Writing – Original Draft Preparation (supporting). All authors read and approved the final manuscript.

## Acknowledgements

This work was supported by the German Network for Bioinformatics Infrastructure-de.NBI, service center BioInfra.Prot, funded by the German Federal Ministry of Education and Research (BMBF)-Grant FKZ 031 A 534A. T. Sauer is grateful for the scholarship he received from the Ad Infinitum Foundation. Additionally, we thank Julia Horn, Katja Klempt-Gießing, and Emma Neumann for their excellent technical assistance in this project.

## Authors’ information

## Notes

### Competing Interest Statement

The authors have declared no competing interest.

## References

1. Ferlay J EM, Lam F, Laversanne M, Colombet M, Mery L, Piñeros M, Znaor A, Soerjomataram I, Bray F. Global Cancer Observatory: Cancer Today (version 1.1). International Agency for Research on Cancer.; 2024 [Available from: https://gco.iarc.who.int/today.

2. Marrugo-Ramírez J, Mir M, Samitier J. Blood-Based Cancer Biomarkers in Liquid Biopsy: A Promising Non-Invasive Alternative to Tissue Biopsy. Int J Mol Sci. 2018;19(10):2877.

3. Lim M, Kim C-J, Sunkara V, Kim M-H, Cho Y-K. Liquid Biopsy in Lung Cancer: Clinical Applications of Circulating Biomarkers (CTCs and ctDNA). Micromachines (Basel). 2018;9(3):100.

4. Drucker E, Krapfenbauer K. Pitfalls and limitations in translation from biomarker discovery to clinical utility in predictive and personalised medicine. EPMA Journal. 2013;4(1):7.

5. Chaltiel D. crosstable: Crosstables for Descriptive Analyses. 2025.

6. Demichev V, Messner CB, Vernardis SI, Lilley KS, Ralser M. DIA-NN: neural networks and interference correction enable deep proteome coverage in high throughput. Nat Methods. 2020;17(1):41–4.

7. Consortium TU. UniProt: the universal protein knowledgebase in 2021. Nucleic Acids Res. 2020;49(D1):D480–D9.

8. Cox J, Hein MY, Luber CA, Paron I, Nagaraj N, Mann M. Accurate proteome-wide label-free quantification by delayed normalization and maximal peptide ratio extraction, termed MaxLFQ. Mol Cell Proteomics. 2014;13(9):2513–26.

9. Zhang X, Smits AH, van Tilburg GB, Ovaa H, Huber W, Vermeulen M. Proteome-wide identification of ubiquitin interactions using UbIA-MS. Nat Protoc. 2018;13(3):530–50.

10. Huber W, von Heydebreck A, Sültmann H, Poustka A, Vingron M. Variance stabilization applied to microarray data calibration and to the quantification of differential expression. Bioinformatics. 2002;18 Suppl 1:S96–104.

11. Gatto L, Gibb S, Rainer J. MSnbase, efficient and elegant R-based processing and visualisation of raw mass spectrometry data. bioRxiv. 2020:2020.04.29.067868.

12. Gatto L, Lilley KS. MSnbase-an R/Bioconductor package for isobaric tagged mass spectrometry data visualization, processing and quantitation. Bioinformatics. 2012;28(2):288–9.

13. Benjamini Y, Hochberg Y. Controlling the False Discovery Rate: A Practical and Powerful Approach to Multiple Testing. Journal of the Royal Statistical Society: Series B (Methodological). 1995;57(1):289–300.

14. Kassambra A. ggpubr: ‘ggplot2’ Based Publication Ready Plots. 0.6.0 ed2023.

15. Pinheiro J, Bates D. Mixed-Effect Models in S and S-plus 2002.

16. R Development Core Team. R: A language and environment for statistical computinng. Vienna, Austria: R Foundation for Statistical Computing; 2021.

17. Simko TWV. R package ‘corrplot’: Visualization of a Correlation Matrix. 0.95 ed 2024.

18. Wickham H, Averick M, Bryan J, Chang W, McGowan L, François R, et al. Welcome to the Tidyverse. Journal of Open Source Software. 2019;4:1686.

19. Kyte J, Doolittle RF. A simple method for displaying the hydropathic character of a protein. J Mol Biol. 1982;157(1):105–32.

20. Guruprasad K, Reddy BV, Pandit MW. Correlation between stability of a protein and its dipeptide composition: a novel approach for predicting in vivo stability of a protein from its primary sequence. Protein Eng. 1990;4(2):155–61.

21. Ikai A. Thermostability and aliphatic index of globular proteins. J Biochem. 1980;88(6):1895–8.

22. Osorio D, Rondón-Villarreal P, Torres Sáez R. Peptides: A Package for Data Mining of Antimicrobial Peptides. The R Journal. 2015;7:4–14.

23. Max Kuhn HW. Tidymodels: a collection of packages for modeling and machine learning using tidyverse principles. 2020.

24. Perez-Riverol Y, Csordas A, Bai J, Bernal-Llinares M, Hewapathirana S, Kundu DJ, et al. The PRIDE database and related tools and resources in 2019: improving support for quantification data. Nucleic Acids Res. 2019;47(D1):D442-D50.

25. Zolg W. The Proteomic Search for Diagnostic Biomarkers: Lost in Translation? *. Molecular & Cellular Proteomics. 2006;5(10):1720–6.

26. Batis N, Brooks JM, Payne K, Sharma N, Nankivell P, Mehanna H. Lack of predictive tools for conventional and targeted cancer therapy: Barriers to biomarker development and clinical translation. Advanced Drug Delivery Reviews. 2021;176:113854.

27. Dill KA. Dominant forces in protein folding. Biochemistry. 1990;29(31):7133–55.

28. Kauzmann W. Some factors in the interpretation of protein denaturation. Adv Protein Chem. 1959;14:1–63.

29. Nicholls A, Sharp KA, Honig B. Protein folding and association: Insights from the interfacial and thermodynamic properties of hydrocarbons. Proteins: Structure, Function, and Bioinformatics. 1991;11(4):281–96.

30. Dias CL, Ala-Nissila T, Wong-ekkabut J, Vattulainen I, Grant M, Karttunen M. The hydrophobic effect and its role in cold denaturation. Cryobiology. 2010;60(1):91–9.

31. Dias CL, Ala-Nissila T, Karttunen M, Vattulainen I, Grant M. Microscopic Mechanism for Cold Denaturation. Physical Review Letters. 2008;100(11):118101.

32. Bhatnagar BS, Pikal MJ, Bogner RH. Study of the Individual Contributions of Ice Formation and Freeze-Concentration on Isothermal Stability of Lactate Dehydrogenase during Freezing. Journal of Pharmaceutical Sciences. 2008;97(2):798–814.

33. Kueltzo LA, Wang Wei, Randolph TW, Carpenter JF. Effects of Solution Conditions, Processing Parameters, and Container Materials on Aggregation of a Monoclonal Antibody during Freeze-Thawing. Journal of Pharmaceutical Sciences. 2008;97(5):1801–12.

34. Mazur P. Cryobiology: the freezing of biological systems. Science. 1970;168(3934):939–49.

35. Mitchell DE, Fayter AER, Deller RC, Hasan M, Gutierrez-Marcos J, Gibson MI. Ice-recrystallization inhibiting polymers protect proteins against freeze-stress and enable glycerol-free cryostorage. Mater Horiz. 2019;6(2):364–8.

36. Strambini GB, Gonnelli M. Protein Stability in Ice. Biophysical Journal. 2007;92(6):2131–8.

37. Hillgren A, Lindgren J, Aldén M. Protection mechanism of Tween 80 during freeze–thawing of a model protein, LDH. International Journal of Pharmaceutics. 2002;237(1):57–69.

38. Krielgaard L, Jones LS, Randolph TW, Frokjaer S, Flink JM, Manning MC, et al. Effect of tween 20 on freeze-thawing-and agitation-induced aggregation of recombinant human factor XIII. Journal of Pharmaceutical Sciences. 1998;87(12):1597–603.

39. Naicker MC, Kim Y-H, Lee K, Im H. Yeast Cyclophilins Prevent Cold Denaturation of Proteins. Bulletin of the Korean Chemical Society. 2016;37(3):366–71.

40. Sanfelice D, Temussi PA. Cold denaturation as a tool to measure protein stability. Biophysical Chemistry. 2016;208:4–8.

41. Bielli P, Calabrese L. Structure to function relationships in ceruloplasmin: a ‘moonlighting’ protein. Cell Mol Life Sci. 2002;59(9):1413–27.

42. Hellman NE, Gitlin JD. Ceruloplasmin metabolism and function. Annu Rev Nutr. 2002;22:439–58.

43. Gitlin JD. Transcriptional regulation of ceruloplasmin gene expression during inflammation. J Biol Chem. 1988;263(13):6281–7.

44. Chen F, Han B, Meng Y, Han Y, Liu B, Zhang B, et al. Ceruloplasmin correlates with immune infiltration and serves as a prognostic biomarker in breast cancer. Aging (Albany NY). 2021;13(16):20438–67.

45. Jia M, Dong T, Cheng Y, Rong F, Zhang J, Lv W, et al. Ceruloplasmin is associated with the infiltration of immune cells and acts as a prognostic biomarker in patients suffering from glioma. Frontiers in Pharmacology. 2023;14.

46. Woong Han I, Jang J-Y, Kwon W, Park T, Kim Y, Bun Lee K, et al. Ceruloplasmin as a prognostic marker in patients with bile duct cancer. Oncotarget. 2017;8(17).

47. Hanas JS, Hocker JR, Cheung JY, Larabee JL, Lerner MR, Lightfoot SA, et al. Biomarker Identification in Human Pancreatic Cancer Sera. Pancreas. 2008;36(1).

48. Teknos TN, Islam M, Arenberg DA, Pan Q, Carskadon SL, Abarbanell AM, et al. The effect of tetrathiomolybdate on cytokine expression, angiogenesis, and tumor growth in squamous cell carcinoma of the head and neck. Arch Otolaryngol Head Neck Surg. 2005;131(3):204–11.

49. Lefranc MP. Immunoglobulin and T Cell Receptor Genes: IMGT(®) and the Birth and Rise of Immunoinformatics. Front Immunol. 2014;5:22.

