## Additional File 2 for "Impact of Single Freeze-Thaw Cycles on Serum Protein Stability: Implications for Clinical Biomarker Validation Using Mass Spectrometry"

T.Sauer et al.

### Additional File 2

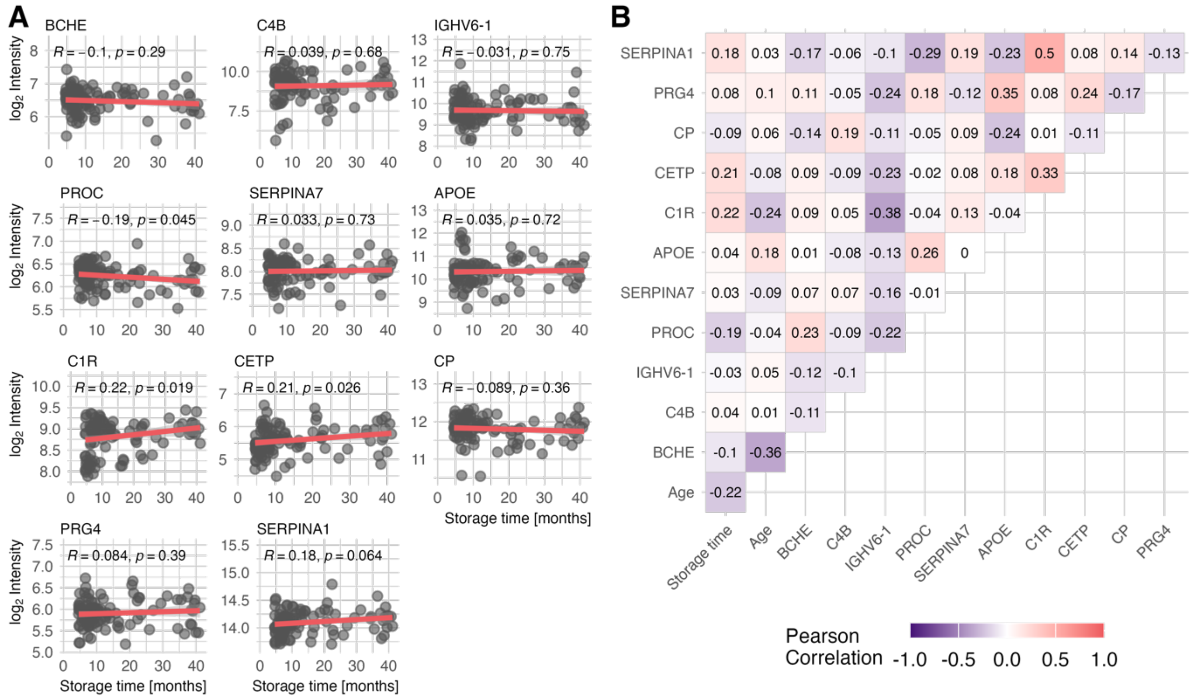


*Supplementary Figure 1: Correlation between protein abundance and storage time in FT samples.* ***A:*** *Scatter plots of $\log_2$ protein intensity and liquid nitrogen storage time in months with linear model trendline and Pearson correlation coefficient. No meaningful correlation was found between protein abundance and storage time of the serum samples.* ***B:*** *No correlation was found between the target proteins and storage time or patient age. Age was included as a negative control variable.*
